# Electrophysiological properties of neurons in the intermediate thoracolumbar spinal cord mediating proprioception

**DOI:** 10.1101/2022.06.23.497422

**Authors:** Felipe Espinosa, Iliodora V. Pop, Helen C. Lai

## Abstract

Animals must know where their body is in relationship to their environment to move appropriately. Proprioception, the sense of limb and body position, is required to produce feedforward predictions of future movement as well as provide feedback information during movement to update the system. How proprioception is processed at the level of the spinal cord is not well understood. Most work has focused on Clarke’s column (CC) neurons, a component of the dorsal spinocerebellar tract (DSCT). CC neurons receive hindlimb proprioceptive information and send it directly to the cerebellum with no axon collaterals to other neurons in the spinal cord. In contrast to CC neurons, we previously found that a subset of thoracolumbar *Atoh1*-lineage neurons also receives proprioceptive afferent inputs, but project locally within the spinal cord including synapsing on motor neurons. We set out to understand how CC and *Atoh1*-lineage neurons in the thoracolumbar spinal cord respond to electrical activity. Using *in vitro* acute spinal cord slices and whole-cell patch clamp, we characterized the passive and active electrical properties of CC and thoracolumbar *Atoh1*-lineage neurons. We find that most CC neurons increase their action potential firing frequency linearly upon increased current injection up to high frequencies likely due to a hyperpolarization-activated current (I_h_). In contrast, most *Atoh1*-lineage neurons do not have an I_h_ current and therefore, their firing frequency fades with increasing current injection. Interestingly, although *Atoh1*-lineage neurons are classically subdivided into a contralaterally-projecting Medial and ipsilaterally-projecting Lateral population, we find no obvious electrophysiological signatures that could distinguish these spatially distinct populations. Finally, we set out to determine the organization of the inputs to CC and *Atoh1*-lineage neurons. In a hemisected spinal cord preparation, while recording from neurons in the lower thoracic to upper lumbar segments, we find that CC neurons receive low threshold inputs (proprioceptive or low threshold mechanoreceptor) from lumbar 2 to 4 dorsal roots, while *Atoh1*-lineage Medial neurons do not. This raises the possibility that *Atoh1*-lineage Medial neurons receive proprioceptive inputs from more local dorsal roots. Altogether, we find that CC and *Atoh1*-lineage neurons have distinct membrane properties and sensory input organization even though they are both reported to receive proprioceptive information and are in close proximity in lamina V-VII of the thoracolumbar spinal cord.

## Introduction

In our daily lives, we move through our environment without thinking about it. Whether reaching for a cup of coffee or scratching one’s back, we have an unconscious sense of the spatial location of each of our body parts. This ‘sixth sense,’ called proprioception, originates from proprioceptive sensory neurons that detect changes in muscle length and tension through specialized receptors called muscle spindles and Golgi tendon organs in the muscle and tendon (Sherrington, 1906; Bosco and Poppele, 2001; Tuthill and Azim, 2018). Alterations in proprioceptive function results in severe locomotor deficits indicating how critical the sense of body position is to motor function (Abelew et al., 2000; Akay et al., 2014; Chesler et al., 2016). How proprioceptive information is received by specific neurons in the spinal cord is not well understood.

Historically, most of the research on spinal proprioceptive circuits has been focused on a prominent set of large neurons in the medial part of the thoracic to upper lumbar spinal cord called Clarke’s column (CC). CC neurons receive hindlimb proprioceptive information and then relay this information directly to the cerebellum through the ipsilaterally-projecting dorsal spinocerebellar tract (DSCT) (Oscarsson, 1965; Bosco and Poppele, 2001; Hantman and Jessell, 2010). Research in cats helped refine our knowledge about neuronal function and connectivity of neurons contributing to the DSCT. Specially, recordings of DSCT neurons using either metal electrodes or glass sharp electrodes in cats under anesthesia or in spinal animals defined responses of DSCT neurons to electrical or physiological stimulation of sensory afferents (Eccles et al., 1961; Oscarsson, 1965; Mann, 1971; Kuno et al., 1973; Edgley and Jankowska, 1988; Bosco and Poppele, 2001). These experiments found cells of the DSCT that responded either to cutaneous touch stimuli, proprioceptive stimuli, or both from the hindlimb. However, CC neurons make up a subset of the DSCT and recordings with sharp and metal electrodes were limited in their ability to assess the cellular response properties of CC neurons (Matsushita and Hosoya, 1979; Baek et al., 2019; Pop et al., 2022). With the advent of patch clamp technology in the 1980s (Hamill et al., 1981), together with current genetic and molecular biology tools, we can precisely identify CC neurons to understand their physiological currents.

In addition to CC neurons, our group discovered that a subset of neurons derived from progenitor cells defined by expression of the basic helix-loop-helix transcription factor, *Atonal homolog 1* (*Atoh1*) also receive proprioceptive information (Yuengert et al., 2015; Pop et al., 2022). CC and *Atoh1*-lineage neurons all reside in laminae V-VII of the thoracolumbar spinal cord. However, in contrast to the long range cerebellar-projecting CC neurons, thoracolumbar *Atoh1*-lineage neurons project locally within the spinal cord including synapsing on motor neurons. Therefore, *Atoh1*-lineage neurons are more integrated into local spinal circuits. In addition, *Atoh1*-lineage neurons are a heterogeneous group. *Atoh1*-lineage neurons cluster into those near the central canal underneath CC neurons (*Atoh1* Medial cells) or lateral to CC neurons (*Atoh1* Lateral cells) (Bermingham et al., 2001; Wilson et al., 2008). *Atoh1* Medial cells, also known as dI1c, are thought to be contralaterally-projecting while *Atoh1* Lateral cells, also known as dI1i, are ipsilaterally-projecting. However, recent evidence suggests that subsets of the *Atoh1* Lateral are also contralaterally-projecting (Kaneyama and Shirasaki, 2018; Pop et al., 2022). Therefore, using whole cell patch clamp and molecular genetic tools in mice, we set out to determine if *Atoh1*-lineage Medial and Lateral neurons had distinguishing electrophysiological signatures.

To further our understanding of how CC and *Atoh1*-lineage Medial and Lateral cells respond to electrical activity and sensory processing, we used modern molecular genetic tools to reproducibly label distinct cell types and a reductionist electrophysiological approach of dissected *in vitro* spinal cord preparations with whole cell patch clamp recordings. Overall, we found that CC neurons are large cells and upon current injection show capacity of firing at higher rates than *Atoh1*-lineage neurons likely due to the presence of a hyperpolarization-activated current (I_h_-like) and related sag potential. We also find that *Atoh1*-lineage Medial and Lateral cells have very similar electrophysiological properties despite their differences in spatial distribution and anatomical projections. Additionally, in line with previous reports, we find that 69% of CC neurons receive low threshold-activated monosynaptic inputs from the lumbar 2-4 (L2-4) roots or a multi-discharge input pattern from sensory afferents, perhaps from converging inputs from several muscles (Holmqvist et al., 1956; Osborn and Poppele, 1988). In contrast, very few *Atoh1*-lineage Medial neurons receive monosynaptic inputs from L2-4 and only 23% of them showed primary afferent connectivity in our experimental conditions with almost all receiving a multi-discharge pattern. Altogether, we find that although CC and *Atoh1*-lineage neurons reside together in lamina V-VII of the thoracolumbar spinal cord, they have unique intrinsic electrophysiological properties and input organization.

## Materials & Methods

### Mouse strains

All animal experiments were approved by the Institutional Animal Care and Use Committee at UT Southwestern. The following mouse strains were used: *Gdnf*^*IRES2-CreERT2*^ (Cebrian et al., 2014)(JAX #024948), *Atoh1*^*Cre*^ knock-in (Yang et al., 2010), and *R26*^*LSL-tdTom*^ (Ai14)(JAX #007914)(Madisen et al., 2010). *Gdnf*^*IRES2-CreERT2*^ or *Atoh1*^*Cre*^ mice were crossed to *R26*^*LSL-tdTom*^ mice to label CC and *Atoh1*-lineage neurons with tdTomato (*Gdnf*^*TOM*^ and *Atoh1*^*TOM*^, respectively). *Gdnf*^*TOM*^ pups (aged P7 and/or P8) were injected with 10 mg/mL tamoxifen (Sigma) dissolved in sunflower oil (Sigma) with 10% ethanol at a dosage of 0.1 mg/g pup. All mice were outbred and thus, are mixed strains (at least C57Bl/6J, C57Bl/6N, and ICR).

### Electrophysiological Recordings

#### Transverse Slices

Mice P10-17 were anesthetized with isoflurane (drop technique) and decapitated. A spinal cord section spanning thoracic and lumbar roots T2-L2 was dissected out by performing dorsal and ventral laminectomies using chilled and oxygenated dissecting solution. The dura mater was cut on the midline and dorsal and ventral roots were cut close to the spinal cord. The isolated spinal cord section was then embedded in 2.5% agarose. Afterwards, dissecting solution was then poured in the dissecting chamber and oxygenated with 95% O_2_/5% CO_2_. Spinal cord slices of 300-350 µm thickness were sliced using a Leica Vibratome 1000 Plus. Transverse sections were obtained between T9-L1. The dissecting solution contained (in mM): 81 NaCl, 26 NaHCO_3_, 75 sucrose, 2.5 KCl, 1.25 NaH_2_PO_4_, 0.5 CaCl_2_, 5 MgCl_2_, and 20 dextrose (305-310 mOsm). Afterwards, slices were incubated in a slightly modified aCSF solution containing (in mM): 130 NaCl, 2.5 KCl, 1.25 NaH_2_PO_4_, 0.75 mM CaCl_2_, 2.5 mM MgCl_2_, 26 NaHCO_3_, and 10 dextrose for 15 minutes at 34°C and then for at least 45 min at room temperature until transferred to the recording chamber.

For transverse slice recordings, the aCSF bath solution contained (in mM): 130 NaCl, 2.5 KCl, 1.25 NaH_2_PO_4_, 2 CaCl_2_, 1.25 MgCl_2_, 26 NaHCO_3_, and 10 dextrose. Whole-cell patch clamp was performed using borosilicate capillaries pulled with a P-87 flaming-brown micropipette puller (Sutter Instruments). The pipettes had a resistance of 3-6 MΩ, when using an internal solution containing (in mM): 134 KCl, 10 NaCl, 1 CaCl_2_, 1 MgCl_2_, 1 dextrose, 4 Na_2_ATP, and 10 HEPES-K (pH 7.25, 290-295 mOsm). A junction potential of 11 mV was determined empirically and was consistent with that calculated theoretically by using the corresponding function in Clampfit. This value was used to adjust voltages reported herein. Recordings were obtained using a 700B Multiclamp Amplifier (Molecular Devices). Neurons were visualized using a Nikon EF600N Eclipse Microscope equipped with infrared differential interference contrast, a CCD camera and epifluorescence. A fluorescence filter set Y-2E/C (Texas Red, EX 540-580, DM 595, BA 600-660) was used to visualize tdTomato labeled neurons. Acquisition was done at 10 kHz and data was analyzed offline using Clampfit Analysis Suite (Molecular Devices), GraphPad Prism 9, and Microsoft Excel 2015.

After achieving whole-cell configuration, cells were held near resting potential (Vm_rest_) for at least 5 minutes to allow for dialysis of the pipette internal solution. Afterwards, when needed, cells were held at −70 mV by injecting current of the appropriate sign and amplitude. Depending on the experiment, one or several of the following protocols were applied: For rheobase evaluation, first, a gross determination of action potential firing was made using 10-20 pA incremental steps (1 s duration), and then a baseline subthreshold current with incremental steps of 1-3 pA in successive sweeps were used. The rheobase was considered the step at where the first action potential (AP) was fired and where APs were consistently fired in the following steps. To determine firing properties and firing types, ten 50 pA/1s current steps were applied. Neurons were classified as pertaining to 1 out of 4 firing-type groups: Tonic (T), Fading (F), Single (S) or Undefined (U). U cells were classified a such because their firing pattern changed depending on the intensity of the stimulus. T-firing type cells showed increasing firing frequency and stable AP amplitude all along the ten steps. Cells were classified as F-firing type if the amplitude of three of the last APs fired in a step were reduced 25% or more compared to the first AP. The F-firing type also presented with AP failure during a sweep, from which they could not recover (i.e. they did not resume firing APs). F-firing type cells also showed a rise in the baseline V_m_ in subsequent APs. To determine the presence of hyperpolarization-activated currents (I_h_) in voltage clamp, and of sag potentials and rebound AP firing in current clamp, cells were held at −70 mV and hyperpolarizing potentials (nine −10 mV steps/1 s), or hyperpolarization currents (up to 20 steps of −50 pA/200 ms) were applied.

#### Spinal Cord Hemisections

A spinal cord section from T2-L6 was dissected out as described for transverse slices. Dorsal roots L2-L4 were preserved for afference stimulation, and 1 dorsal root from T11 to T13 was preserved as anatomical reference. The dura mater was cut on the midline with scissors, and the arachnoid and the pia mater were cut using a 21-gauge needle. Left and right spinal cord sides were then separated to obtain hemisections. After incubation in the modified aCSF solution for 15-30 min at 34°C, hemisections were placed in a submersion-recording chamber with the midline facing up.

Recordings were then performed in oxygenated aCSF superfused at a rate of 2-3 ml/min and supplemented with 100 µM picrotoxin (Alomone Labs) and 0.5-1 µM Strychnine (Sigma). The internal solution was the same as above with 2-4 mM QX314 to block Na_+_ channels from driving AP firing. In this recording configuration, only CC neurons in the *Gdnf*^*TOM*^ mouse model, or Medial neurons in the *Atoh1*^*TOM*^ mouse model were at depths suitable for recordings. Lateral cells are too deep to be recorded using this approach. Recordings were obtained between segments T10-L1 by performing whole-cell patch-clamp. Suction electrodes (A-M Systems) were used to stimulate the lumbar dorsal roots (L2-L4). Fluorescently (tdTomato) labeled cells were recorded in a whole-cell configuration one or more segments rostral to the stimulated root. Using a A365 stimulus isolator (WPI), 0.2 ms electric shocks of increasing magnitude were applied starting around 5 µA and up to 500 µA. The stimulus threshold was determined by gradually increasing the stimulation intensity until at least 3 out of 5 stimulation attempts yielded a synaptic response. To target proprioceptive and low-threshold mechanoreceptor inputs, cells with a threshold below 15 µA were stimulated using a 25 µA stimulus. For those with thresholds between 15 µA-40 µA, a 1.5-2X threshold stimulation was used, but no higher than 60 µA. Cells with thresholds above 40 µA were not further analyzed.

### Experimental Design and Statistical Tests

Raw traces were analyzed using Clampfit (Molecular Devices). All data and graphs were processed in Microsoft Excel 2015 and GraphPad Prism 9. Mean ± SEM are reported throughout the manuscript. *P* values are included in the figures. Statistical tests used are detailed in the Results and/or Figure Legends.

## Results

### Distinct passive and active properties of Clarke’s Column neurons (CC) and Atoh1-lineage Medial and Lateral neurons

CCs and *Atoh1*-lineage cells both reside within laminae V-VII of the spinal cord and are reported to receive proprioceptive inputs (Fig. 1A) (Hantman and Jessell, 2010; Yuengert et al., 2015). We focused on a comprehensive electrophysiological characterization of CC, *Atoh1*-lineage Medial, and *Atoh1*-lineage Lateral cell groups utilizing the whole cell approach in patch clamp experiments. Their basic excitability properties were determined in transverse slices 300-350 µm thick. To test their baseline activity, no holding current was applied so that the cell was recorded at its resting membrane potential, Vm_rest_. Examples of spontaneously firing cells at Vm_rest_ and of the rheobase determination are shown (Fig. 1B, upper panels and lower panels, respectively). CC neurons show a 4-to 8-fold larger capacitance than *Atoh1*-lineage Medial and Lateral cells (CC, 138.1 ± 0.11pF, n=36, Medial, 19.35 ± 1.95 pF, n=23, and Lateral 30.9 ± 2.90 pF, n=30. Welch’s t-tests: CC vs. Medial, *P* < 0.0001, CC vs Lateral, *P* < 0.0001, and Medial vs. Lateral 0.0025). We also find differences in input resistance (Rm)(CC, 0.41 ± 0.10 GΩ, n=38, Medial, 1.81 ± 0.31 GΩ, n=24, and Lateral 0.99 ± 0.212 GΩ, n=29. Welch’s tests: CC vs. Medial, *P* = 0.0004, CC vs Lateral, *P* = 0.0163, and Medial vs. lateral 0.0426) and resting potential (Vm_rest_)(CC, −58.91 ± 1.43 mV, n=35, Medial, −53.75 ± 0.98 mV, n=24, and Lateral −51.9 ± 1.72mV, n=29. Welch’s tests: CC vs. Medial, *P* = 0.0045, CC vs Lateral, *P* = 0.0028, and Medial vs. Lateral, ns)(Fig. 1C). Roughly 45% of CC cells spontaneously fire action potentials (APs) at rest while 65% and 40% of *Atoh1*-lineage Medial and Lateral cells, respectively, fire spontaneously. No differences in spontaneous firing frequency were found (Fig. 1C).

**Figure 1.**
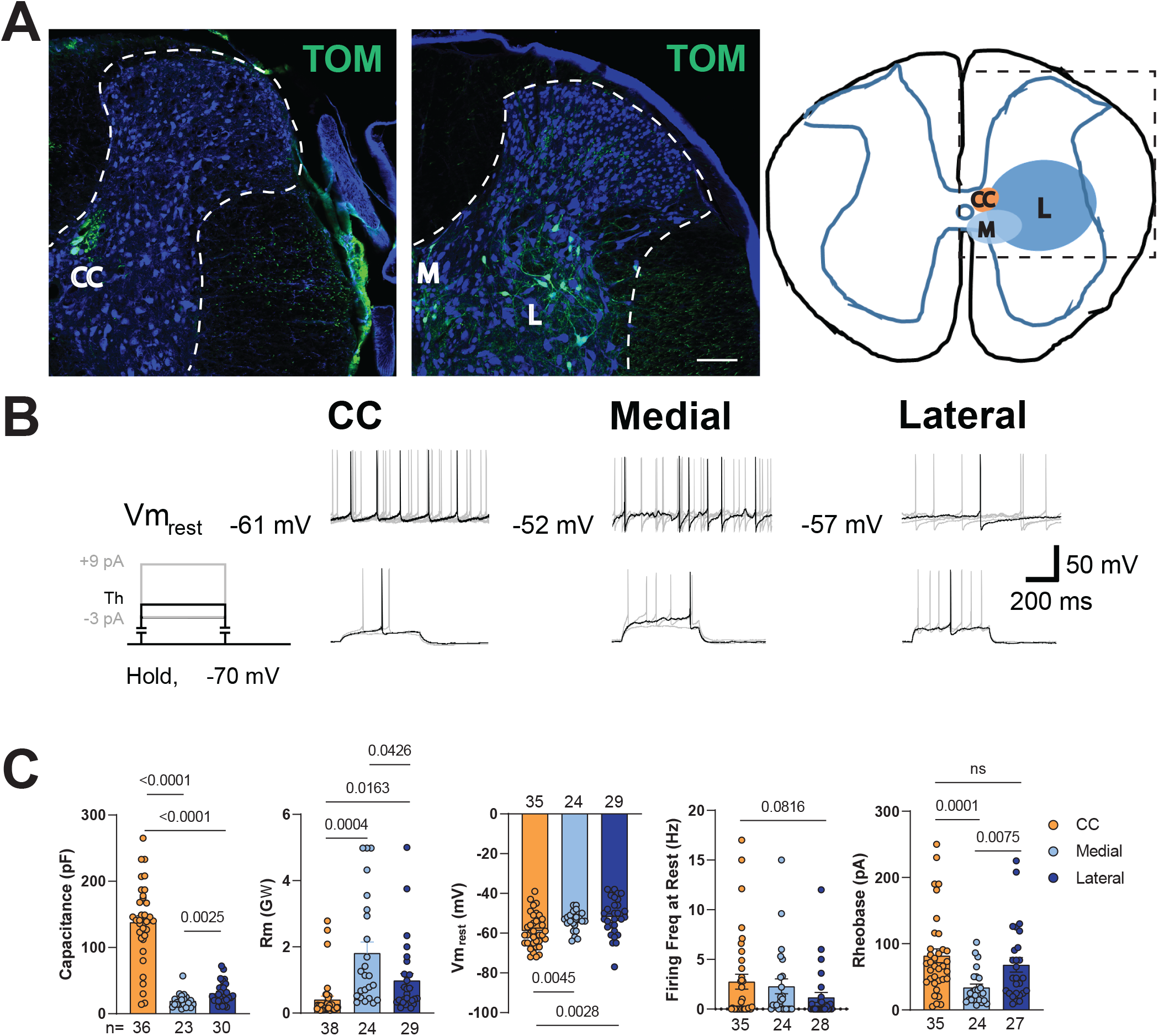
Electrophysiological properties of CC and *Atoh1*-lineage Medial and Lateral neurons. A. Representative lower thoracic images of CC neurons in *Gdnf*^*TOM*^ mice and Medial (M) and Lateral (L) cells in *Atoh1*^*TOM*^ mice. TOM is pseudocolored green. Scale bar 100 µm. All CC and *Atoh1*-lineage M and L cells are within laminae V-VII of the intermediate spinal cord. **B. Top Row**. Some cells fire APs at their resting membrane potential Vm_rest_ (i.e. no holding currents). For each sample trace, Vm_rest_ is shown to the left. There are 5 traces corresponding to five 2 sec sweeps, where only one trace is ‘highlighted’ in black, the rest are shown as background in light gray. **Bottom Row**. The protocol schematic to the left indicates that cells were held at −70 mV. After an initial estimation of the rheobase, a more precise determination was done by current steps of 1-3 pA where the first step was a few pAs below the initially estimated threshold. The schematic and traces only show the sweep at threshold in black and, in light gray, sweeps 3 pA below and 9 pA above threshold. **C**. Quantification for capacitance, input resistance (Rm), Vm_rest_, firing frequency at rest, and rheobase.

To test for excitability, we determined the rheobase. Consistent with the difference in capacitance and internal resistance, CC and *Atoh1*-lineage Lateral neurons show larger rheobase than *Atoh1*-lineage Medial cells (CC, 81.83 ± 10.4 pA, n=35, Medial, 32.6 ± 4.79 pA, n=24, and Lateral 68.15 ± 11.15 pA, n=27. Welch’s tests: CC vs. Medial, *P* = 0.0001, CC vs Lateral, *ns*, and Medial vs. Lateral, 0.0075)(Fig. 1C). Next, we determined the firing properties of these neurons subjected to discrete increases in intracellular current injections. Upon 10 pulse steps at 50 pA increments, the firing properties were classified in either of 4 groups: tonic (T), fading (F), single (S), or undefined (U) (see Materials and Methods for classification criteria). The T-firing type was stable across the stimulation spectrum with little to no accommodation. CC T-firing type cells respond linearly to current injection (Fig. 2C). In contrast, the baseline became clearly depolarized in the F-firing type the larger the amount of current injected (Fig. 2A), resulting in eventual firing failure during the 1 s step. This phenomenon may be due in part to depolarization block where suboptimal repolarization results in a slower recovery from inactivation of fast voltage-dependent Na_+_-channels (Grace and Bunney, 1986; Richards et al., 1997; Qian et al., 2014). CC and *Atoh1*-lineage Medial and Lateral F-firing type cells had different kinetics of AP firing upon current injection (Fig. 2C, Fading). CC F-firing type cells reached a plateau around 450 pA, while *Atoh1*-lineage Medial F-firing type cells reached a plateau around 100 pA and Lateral F-firing type showed an inverted “U” shape (Fig. 2C). The S-firing type was defined as cells that had less than 3 APs at all current injections. The S-firing type seems an extreme case of F-type firing, where the decrease in AP amplitude is very steep and firing fails quickly (dashed blue lines, Fig. 2A).

**Figure 2.**
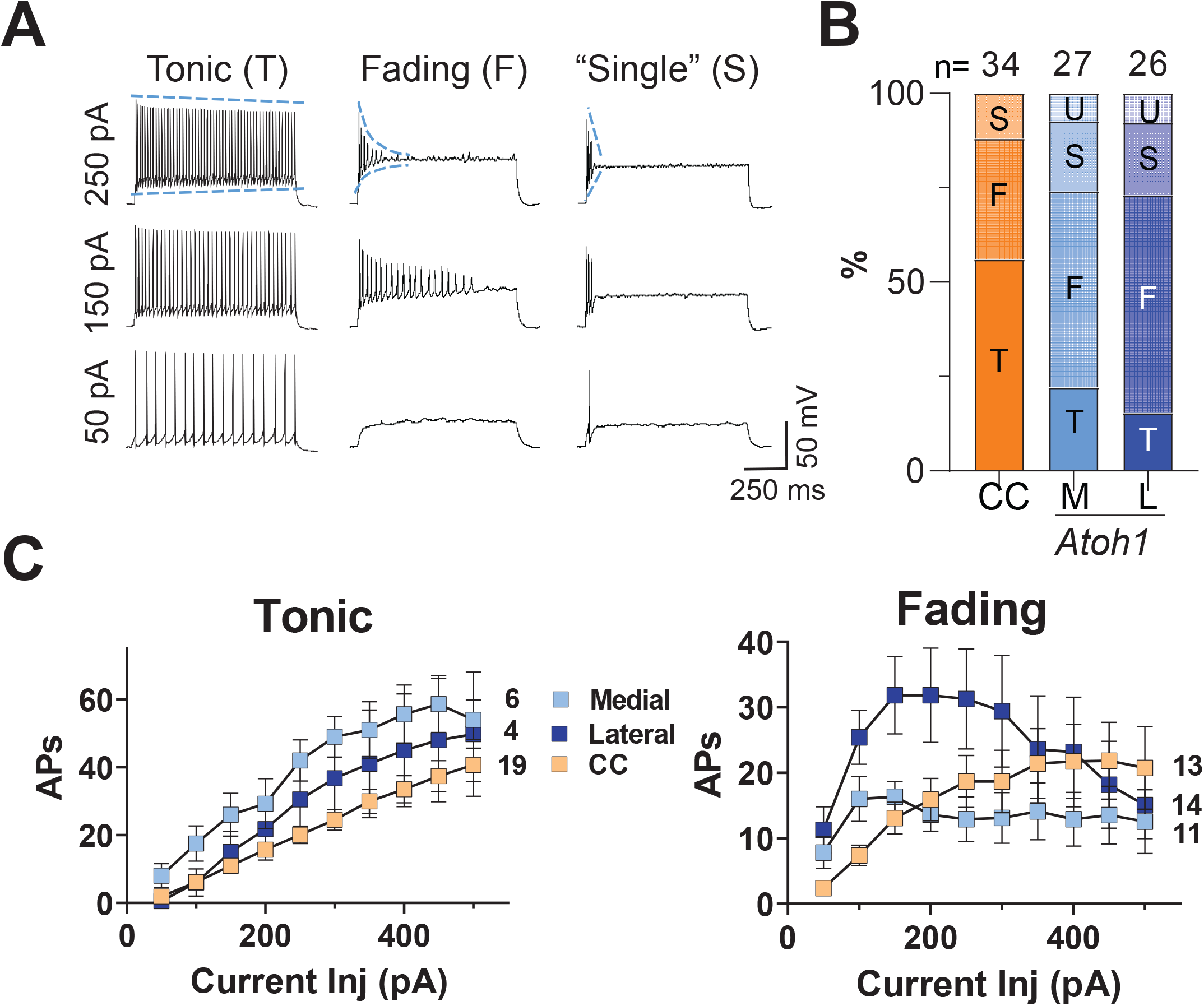
Firing types for CC and *Atoh1*-lineage Medial and Lateral neurons. **A**. Current steps in increments of 50 pA were injected up to 500 pA. Steps at 50, 150 and 250 pA are shown. Columns show the main firing types found for all cell groups. The Tonic-firing type (T) showed a linear increase in frequency at higher steps with no failures. The Fading-firing type (F) showed increased firing frequency upon the first few depolarization steps but with further depolarization failed to sustain firing. The ‘Single’-firing type (S) showed less than 3 APs, even at the highest currents injected. The blue lines on the top traces are shown to highlight the slope differences among the firing types for the decrease in maximal AP amplitude and in baseline depolarization. The slope is shallow for tonic firing but very steep for fading and single firing cells. **B**. Percentages of firing types for each cell group. M and L neurons that had cells that changed firing type depending on the level of current injected were classified as undefined (U). **C**. The relation for current injected and # of APs fired is shown for T- and F-firing types for each cell group.

The proportion of firing-types differs among groups (Fig. 2B). T-firing type is predominant in CC cells, whereas it is lower in Medial and Lateral *Atoh1*-lineage cells (CC, 56%, Medial 22.2%, and Lateral 15.4%). In contrast, the F-firing type is more common in *Atoh1*-lineage cells (CC, 32%, Medial 51.9%, and Lateral 57.7%). The S-firing type is present in lower numbers in all the groups (CC, 12%, Medial 18.5%, and Lateral 19.2%). Interestingly, some cells in Medial and Lateral groups switched among different firing-types depending on the strength of the stimulation and were categorized as U cells. Similarly, it has previously been shown that some neurons can switch from tonic firing to burst firing under the influence of neuromodulators (Grace and Bunney, 1984a, b; Zhang, 2003), suggesting that firing-type classification is not absolute and may better be interpreted as the predominant physiological state of a neuron under those experimental conditions and developmental stage.

### Hyperpolarization-activated- (I_h_) like currents and sag potentials are present in CC cells

When subjected to hyperpolarizing voltage or current steps, CC cells present the activation of a current that resembles I_h_ in voltage clamp and sag potentials in current clamp (Fig. 3A). We injected hyperpolarizing current from a baseline of −70 mV (Fig. 3A). The rate of activation and amplitude of I_h_-like currents increased with each voltage step. The I_h_-like current increases with increasing hyperpolarization (Fig. 3A, inset on left side, −118 ± 20.6 pA at Δ-30 mV, −408 ± 50.3 pA at Δ-60 mV, and −616 ± 81.5 pA at Δ-90 mV, n=10). Similarly, current steps of successive −50 pA increments, induced sag potentials that increased in rate and amplitude with increased hyperpolarization (Fig. 3A, right side). In contrast, *Atoh1*-lineage neurons did not evidence an I_h_ current upon hyperpolarizing voltage or current steps (Fig. 3B). Moreover, *Atoh1*-lineage neurons were more susceptible to damage due to hyperpolarization. If injected with current to more negative than around −120 mV, these cells showed anomalous potential transitions that could resemble irregular step depolarizations (Fig. 3B, right side). Often, the usually high Rm was lost, suggesting plasma membrane leakage and damage.

**Figure 3.**
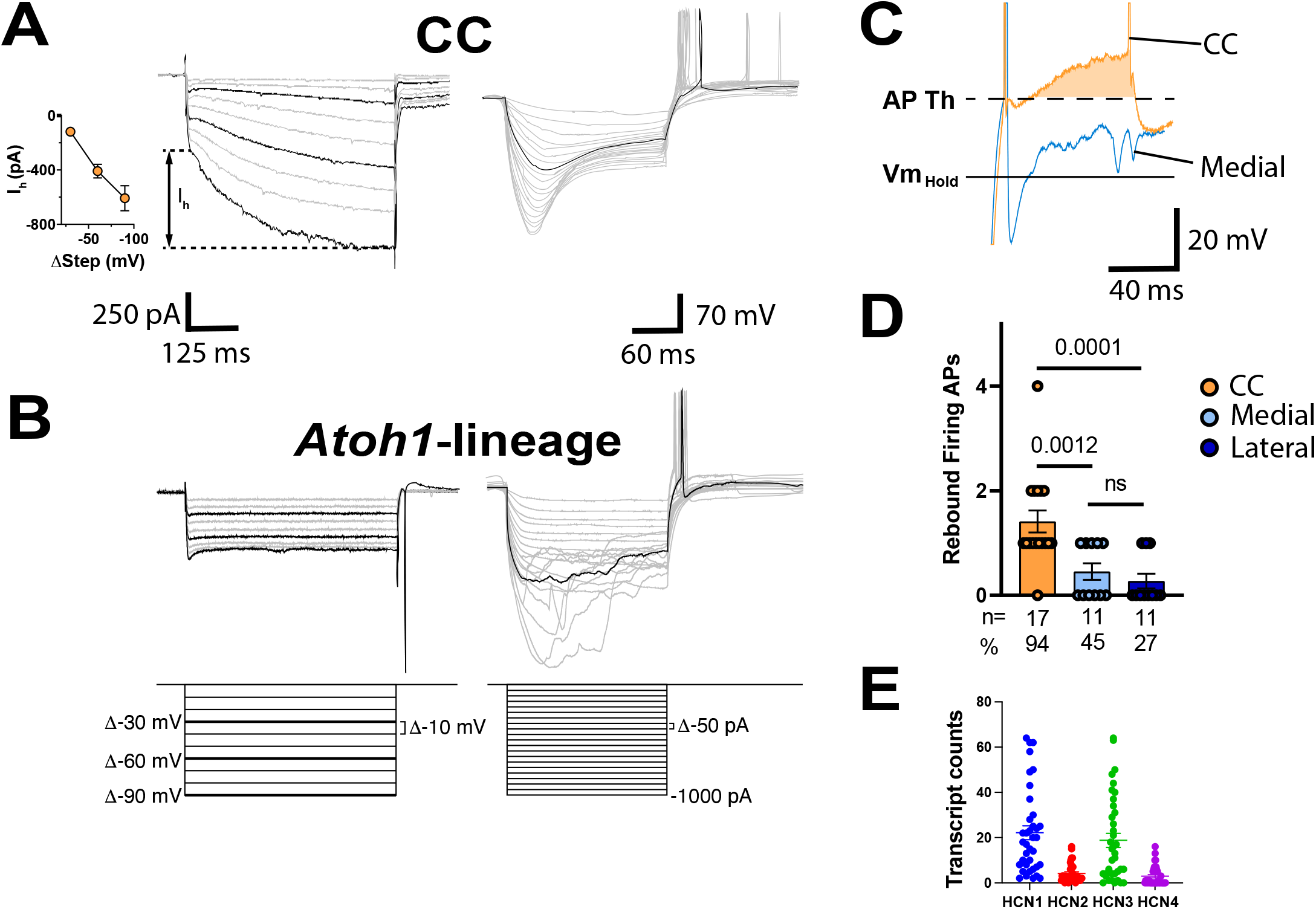
I_h_ currents and sag potentials are present in CC cells. **A**. Hyperpolarizing −10 mV voltage steps (left), and −50 pA current steps (right) induced I_h_-like currents and sag potentials in CC cells. The inset to the left in A shows the quantification for the amplitude for the I_h_-like current induced by steps at −30 mV, −60 mV and −90 mV, after the subtraction of the instantaneous leak current. **B**. Neither I_h_-like currents nor sag potentials were induced in *Atoh1*-lineage neurons. Instead, *Atoh1*-lineage neurons showed high susceptibility to damage when subjected hyperpolarization. As an example, voltage traces to the right became erratic (seen as irregular jumps of the membrane potential) after reaching a certain threshold. **C**. Compared to *Atoh1*-lineage Medial neurons, CC cells showed a relatively large after depolarization (orange-shaded area) following the first rebound AP fired. **D**. The maximal # of APs fired following a hyperpolarization step. **E**. Transcript counts for HCN1, 2, 3, and 4 transcripts in CC neurons from single cell RNA-seq data published in Baek et al.

After hyperpolarization, some cells had rebound firing. A closeup of the rebound firing in a CC cell reveals that the AP is mounted on a transient depolarization that may facilitate repetitive-firing after rebound that is not present in *Atoh1*-lineage neurons (orange shaded area in Fig. 3C). The maximum number of APs fired after rebound was higher (CC, 1.44 ± 0.21 APs, n=17, Medial, 0.455 ± 0.16 pA, n=11, and Lateral 0.27 ± 0.14 pA, n=11. Welch’s tests: CC vs. Medial, *P* = 0.0012, CC vs Lateral, *P* = 0.0001, and Medial vs. Lateral, ns)(Fig. 3D). Rebound firing was more common in CC cells with 94% of CC cells (16/17), 45% of Medial cells (5/11) and 27% of Lateral cells (3/11) showing rebound firing with a hyperpolarization threshold to rebound-firing around - 140 mV (Fig. 3D). *Atoh1*-lineage neurons never had rebound-firing more than 1 AP.

I_h_ is driven by a family of hyperpolarization-activated cyclic-nucleotide gated (HCN) ion channels (Pape and McCormick, 1989; Ludwig et al., 1998). To examine which HCN channels are expressed in CC neurons, we analyzed single cell RNA-sequencing data of spinocerebellar neurons in aged P6-P7 mice from Baek et al. (Baek et al., 2019). Cluster SCT5 in their dataset expresses markers consistent with CC neurons. An analysis of transcript counts in 37 cells of cluster SCT5 found HCN1 and 3 to be enriched (Fig. 3E).

### Low-threshold lumbar afferent inputs to the intermediate spinal cord

CC neurons are considered the main spinal cord cells receiving proprioceptive information. We also reported that some cells within the *Atoh1*-lineage Medial and Lateral populations in the intermediate spinal cord receive proprioceptive inputs (Yuengert et al., 2015). To test whether both CC and *Atoh1*-lineage neurons receive low-threshold inputs from the lumbar spinal cord, we recorded electrically driven afferent signals *in vitro* in hemisected spinal cords. With this sagittal approach, only CC and *Atoh1*-lineage Medial neurons were targeted. We stimulated L2-L4 dorsal roots while recording from CC or *Atoh1-*lineage Medial neurons one to several segments rostral of the dorsal root being stimulated (Fig. 4A). We recorded 62 CC neurons and 92 *Atoh1*-lineage Medial neurons.

**Figure 4.**
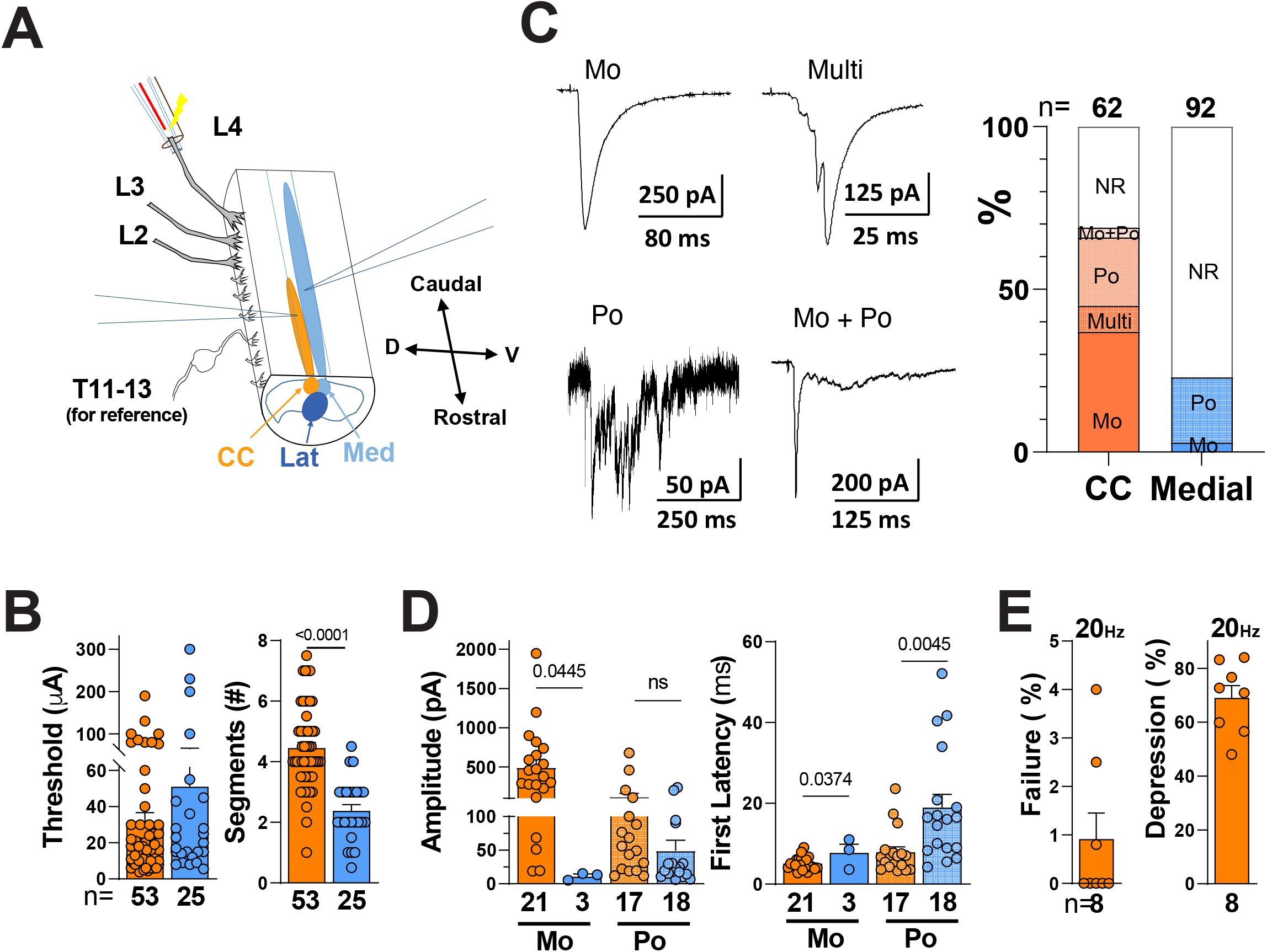
Properties of low-threshold primary afferents for CC and *Atoh1*-lineage Medial neurons in the intermediate spinal cord. **A**. Schematic of electrophysiological recordings in a hemisected spinal cord. Suction electrodes were used to stimulate dorsal roots L2, L3, or L4, while recording from CC or *Atoh1*-lineage Medial neurons in spinal cord segments T10-L1. The estimated location for CC and *Atoh1*-lineage M and L cells is color coded. **B**. The overall threshold stimulation to any response and the number of segments between the recorded cell and stimulated dorsal root are shown. These include cells that responded to any stimulation intensity. **C**. Samples of the four types of EPSCs recorded upon dorsal root stimulation: monosynaptic (Mo), Multi-input (Multi), Polysynaptic (Po) and Mixed (Mo+Po). The representative Po sample was taken from a M cell, all the others are from CC cells. The quantification of EPSC-type percentages for CC and M cells at low-threshold stimulation (< 60 µA) is shown to the right. Cells that did not show response (NR) were also included for the calculation. Mo EPSCs were the most frequent EPSC type found among CC cells, followed by Po EPSCs. Only 23% of M cells showed a response (i.e., 77% were NR) at low-threshold stimulation, with almost all of them showing small Po EPSCs. No Multi or Mo+Po were present in M cells. **D**. Amplitude and First Latency for Mo and Po EPSCs are shown. **E**. Mo EPSCs in CC cells did not show any significant # of failures when stimulated at a frequency of 20 Hz. When amplitudes of the first versus the last EPSC in a train were compared, most of them showed depression above 50%.

First, to get a general sense of the connectivity during our experiments regardless of sensory input type, we determined the threshold to respond to an afferent stimulus from intensities up to 500 µA (CC, 31.6 ± 5.10 µA, n=53; Medial, 51.0 ± 15.33 µA, n=25, *ns*)(Fig. 4B). 85% of CC neurons (53/62 cells) and 27% of Medial cells (25/92 cells) showed a response at any stimulation intensity. CC neurons in the thoracolumbar spinal cord have been shown to receive lumbar and sacral sensory inputs using degeneration and stimulation experiments in cats (Szentagothai and Albert, 1955; Holmqvist et al., 1956; Eccles et al., 1961). Consistent with these results, we found that stimulating the L2-4 dorsal roots resulted in afferent responses in CC cells up to 7 segments rostral to the excited dorsal root (Fig. 4B). Connectivity of CC neurons farther than 7 segments away from the excited dorsal root were not tested. *Atoh1*-lineage neurons could also be triggered rostral to the stimulated dorsal roots, however, the number of segments apart that could trigger a response was less than CC neurons (CC, 4.47 ± 0.18 segments, n =53; Medial, 2.38 ± 0.20 segments, n = 25, *P* = <0.0001)(Fig. 4B).

Upon low threshold stimulation (less than 60 µA) of an L2-L4 dorsal root, we found four different response types in CC neurons and only 2 types in *Atoh1*-lineage Medial neurons (Fig. 4C). The most common afferent signal in CC cells was of the monosynaptic (Mo) type, which had a single evoked excitatory postsynaptic current (eEPSC) upon stimulation of a dorsal root. CC Mo inputs had an average latency under 5 ms (Fig. 4D, right). A subset of these CC Mo inputs were subjected to high frequency stimulation, which showed a small latency variance (not shown) and a low incidence of eEPSC failures (only 0.9 ± 0. 19% failures from a 1s, 20 Hz stimulation, n=8) suggesting these inputs are monosynaptic (Fig. 4E)(Nakatsuka et al., 2000; Doyle and Andresen, 2001; Torsney and MacDermott, 2006). Interestingly, Mo events showed strong amplitude depression between the first and last eEPSCs at 20 Hz stimulation (69 ± 1.65%, n=8). This suggests that even though low-threshold primary afferents can be triggered reliably, they are subject to modulation or to short-term depression. The second response type for CC cells had multiple events (Multi). The third response type had a surge of eEPSCs, which we interpreted to be polysynaptic (Po) inputs. The last response type had a mixture of Mo and Po inputs. For CC neurons, 37% received Mo inputs, 8% Multi, 21% Po, 3% Mo + Po, and 31% were not responsive (NR). For *Atoh1*-lineage Medial neurons, 3% received Mo inputs, 20% Po, and 77% NR)(Fig. 4B). Therefore, upon low-threshold stimulation (less than 60 µA), which includes proprioceptive and low-threshold mechanoreceptive afferents, 69% of CC neurons (43/62 cells) had a response, whereas 23% of Medial neurons (21/92 cells) had a response (Fig. 4C).

The eEPSC amplitude and latencies had some differences for CC and *Atoh1*-lineage Medial neurons. CC Mo inputs had large eEPSC amplitudes compared to the few *Atoh1*-lineage Medial cells that had Mo inputs, while the amplitude for the first eEPSC of Po inputs was similar (Fig. 4D)(absolute amplitude Mo eEPSCs: CC, 491.6 ± 100.2 pA, n=21, Medial, 11.3 ± 3.1 pA, n=3. *P* = 0.0445; absolute amplitude Po EPSCs: CC, 120.5 ± 43.5 pA, n=17, Medial, 49.0 ± 15.9 pA, n=18, *ns*). Given that Po events show multiple peaks that are hard to follow from sweep to sweep, the analysis was focused on the latency to the first event (first latency). The first latency for Mo eEPSCs were slightly longer for Medial cells, while the first latency for Po eEPSCs to *Atoh1*-lineage Medial neurons were about 2-3 fold greater than to CC cells (Fig. 4D)(Mo EPSCs: CC, 5.10 ± 0.35 ms, n=21, Medial, 7.7 ± 2.17 ms, n=3, *P* = 0.0374; Po EPSCs: CC, 7.87 ± 1.70 ms, n=17, Medial, 18.9 ± 3.26 ms, n=18, *P* = 0.0045.)

## Discussion

Here, we characterize the passive and active membrane properties of neurons in the spinal cord that are reported to receive proprioceptive information and reside in lamina V-VII of the thoracolumbar spinal cord. In addition, we assess the synaptic inputs they receive from lumbar dorsal roots. We report unique features of the CC and *Atoh1*-lineage populations.

### Clarke’s column (CC)

CC neurons are a major contributor of the DSCT. Vast physiological literature has shown that DSCT neurons including CC cells are triggered by hindlimb muscle stretch and joint position (Oscarsson, 1965; Mann, 1973; Matsushita and Hosoya, 1979; Bosco and Poppele, 2001; Stecina et al., 2013). Anatomical studies in cats and our work in mice show that CC axons are direct highways to granule cells in the cerebellum with no axon collaterals to the spinal cord, medulla, or cerebellar nuclei (Houchin et al., 1983; Walmsley and Nicol, 1990; Pop et al., 2022). We also found that CC neurons make 43-47% of all spinocerebellar neurons (Pop et al., 2022). Therefore, it is important to understand how this major cell type responds to electrical activity. Using the whole cell patch clamp technique, we found that they have a large capacitance consistent with their large morphology. Interestingly, although we label CC neurons with a genetic mouse line that is anatomically homogenous (Pop et al., 2022), we find that CC neurons are electrophysiologically heterogeneous. 56% CC neurons are of the T-firing type and 32% of them are the F-firing type and 12% of the S-firing type. This electrophysiological heterogeneity may result from the acute transverse slice recording preparation or may represent true differences in CC neuron responses. Recordings from *in vivo* preparations could address these possibilities. Interestingly, we found that ∼45% CC neurons fire spontaneously in our acute slice preparation. This is on the same order as recordings in anesthetized or decerebrate cats that found 34% of CC cells or 40% of DSCT cells fire spontaneously *in vivo* (Zytnicki et al., 1995; Fedirchuk et al., 2013). These data suggest that CC neurons in mice and cats may have similar intrinsic firing properties in both *in vitro* and *in vivo* preparations.

Importantly, CC neurons of the T-firing type increase their firing frequency linearly with increasing current injection consistent with literature in cats that find that AP firing of DSCT neurons follows the frequency stimulation of various hindlimb nerves (Holmqvist et al., 1956). Therefore, most CC neurons are frequency encoders with increasing input from muscle activity linearly increasing action potential firing. CC neurons are reported to fire greater than 500 Hz in cats and with instantaneous frequencies as high as 1 kHz with little adaptation (Holmqvist et al., 1956; Kuno and Miyahara, 1968; Eide et al., 1969). In mice, we tested the ability of CC neurons to follow dorsal root stimulation of inputs up to 20 Hz. With a 500 pA depolarizing current injection for 1 second, the highest frequency reached in CC cells was 109 Hz (30 Hz on average).

In part, the ability of CC neurons to fire tonically at high frequencies could be due to the presence of an I_h_-like current. Present in 94% of CC cells recorded, I_h,_ known as the pacemaker current, is driven by a family of hyperpolarization-activated cyclic-nucleotide gated (HCN) ion channels (Pape and McCormick, 1989; Ludwig et al., 1998). These channels are important in setting the resting membrane potential, signal integration, and serve a pacemaker role in different cell types (DiFrancesco and Ojeda, 1980; McCormick and Huguenard, 1992; Maccaferri et al., 1993; Pape, 1996; Bal and McCormick, 1997; Robinson and Siegelbaum, 2003). Single cell RNA-sequencing of spinocerebellar neurons suggests that HCN1 and HCN3 are expressed in CC neurons (Baek et al., 2019). The I_h_-like current could explain why recordings of DSCT neurons in cats show a “long-lasting repetitive discharge” that continues after a stimulus (Holmqvist et al., 1956; Kuno and Miyahara, 1968; Eide et al., 1969). The I_h_ current is activated at voltages below −40 mV indicating that a proportion of these channels are active at rest. After activation of HCN channels at hyperpolarizing voltages, a depolarizing tail current remains upon returning to baseline potential. This depolarizing tail current likely represents closing HCN channels and temporarily keeps the membrane potential at AP firing threshold levels allowing for repetitive firing. Here we confirm that CC cells show a transient depolarization after the initial rebound AP that may help to sustain repetitive firing (Fig. 3C). Similarly, a “residual” depolarization phenomenon was found in *in vivo* cat experiments while recording from DSCT neurons under electrical stimulation of dorsal roots or sensory stimulation (Kuno and Miyahara, 1968; Eide et al., 1969). In contrast to our *in vitro* mouse experiments, rebound firing was not detected in cats, but it is possible that membrane potential clamping was suboptimal, or that cells were damaged in the cat studies. Other culprits for the repetitive discharge besides the I_h_-like current have been proposed. Among them the activation of a persistent Na_+_-current or the activation of Ca_2+_-currents that keep the cell membrane potential in the range of AP firing threshold. Interestingly, both currents participate in oscillatory activity in other cell types (Bal and McCormick, 1997; Yamada-Hanff and Bean, 2013). However, the characteristic sag current in CC cells suggests that the I_h_ current is from HCN channels. More studies are needed to determine the importance of I_h_ or of other players in repetitive firing in CC cells. In addition, it will be important to investigate if CC cells show oscillatory activity under the influence of neuromodulators or under physiologically relevant conditions.

Finally, 69% of CC neurons recorded in a hemisected spinal cord preparation received monosynaptic or polysynaptic inputs from L2-L4 dorsal roots when stimulated at low threshold. These findings are consistent with the well-established literature of CC neurons relaying hindlimb proprioception (Holmqvist et al., 1956; Eccles et al., 1961). Interestingly, for monosynaptic inputs to CC neurons, the average eEPSC amplitude decreased by 69% between the first and last eEPSCs at 20 Hz stimulation for 1 second, suggesting that there is some modulation of the eEPSC. The depression is not due to inhibitory currents because these experiments were performed with inhibitory current blockers. Future experiments are needed to determine the cause of this eEPSC depression perhaps due to neuromodulators, short-term depression or changes in intrinsic currents.

### Atoh1-lineage neurons

In contrast to CC neurons, to the best of our knowledge, this is the first report of the electrical activity of *Atoh1*-lineage neurons. *Atoh1*-lineage neurons have classically been subdivided into a contralaterally-projecting Medial (dI1c) and ipsilaterally-projecting Lateral (dI1i) population (Bermingham et al., 2001; Wilson et al., 2008). However, recent anatomical studies challenge this view with at least a subset of *Atoh1*-lineage Lateral cells projecting potentially both ipsi- and contralaterally (Kaneyama and Shirasaki, 2018; Pop et al., 2022). Without unique molecular markers to distinguish subsets of the *Atoh1*-lineage population, we set out to evaluate whether the Medial and Lateral cells had distinct electrophysiological signatures. While *Atoh1*-lineage Medial and Lateral neurons had some significant differences in capacitance and rheobase, for the most part, they were very similar. Strikingly, even the percentage of firing types in Medial compared to Lateral neurons was very similar with most being the F-firing type. In contrast to CC neurons, *Atoh1*-lineage neurons were much smaller cells as indicated by their capacitance and had less of the T-firing type consistent with the finding that we found no cells with an I_h_ current.

In addition to CC cells, histological and optogenetic evidence showed that other cell types such as the *Atoh1*-lineage neurons can receive low threshold inputs including proprioceptive inputs (Zampieri et al., 2014; Yuengert et al., 2015; Pop et al., 2022). While some *Atoh1*-lineage Medial and Lateral neurons have been found to receive proprioceptive inputs, it is unclear if all of them do. We can detect axo-somatic proprioceptive inputs on *Atoh1*-lineage soma that are nearest the spinal cord-terminating proprioceptive afferents using immunohistochemistry and we can see optogenetically-activated EPSCs in some *Atoh1*-lineage neurons. Interestingly, in this study, we found that thoracolumbar *Atoh1*-lineage Medial neurons did not receive significant monosynaptic low threshold inputs from L2-L4 dorsal roots. Together with our previous data, there are two possibilities for why we see this result. First, while *Atoh1*-lineage Medial neurons do not receive low threshold inputs from L2-L4, they may receive low threshold inputs from dorsal roots closer to the site of recording (i.e. T10-L1). Second, we could only record from the *Atoh1*-lineage Medial neurons that were most superficial to the midline in the hemisected spinal cord preparation. Therefore, it is possible that *Atoh1*-lineage Medial neurons deeper in the tissue receive lumbar low threshold inputs. Future work will test these two possibilities.

The electrophysiological characteristics of CC and *Atoh1*-lineage neurons we report here serve as the foundation for understanding how these cells respond to electrical activity. This information will be valuable for future experiments focused on how these neurons convert sensory signals to motor output.

## Acknowledgements

This work was supported by the Rita Allen Foundation, Welch Foundation I-1999-20190330, Kent Waldrep Foundation, NIH/NINDS R21NS099808, and NIH/NINDS R01NS100741. We thank Lin Gan for the *Atoh1*^*Cre/+*^ knock-in mouse.

